# Deep Learning links TP53 genotype to expression-defined transcriptional program in Acute Myeloid Leukemia

**DOI:** 10.1101/2025.11.10.687690

**Authors:** Miltiadis Tsesmelis, Rebecca Andersson, Alba Ferrer Perez, Sara Montserrat Vazquez, M. Carolina Florian, Medhanie A. Mulaw

## Abstract

Acute myeloid leukemia (AML) is a hematological cancer characterized by genetic diversity and poor clinical outcomes. Among various genetic mutations in AML, TP53 mutations have relatively low prevalence and present very dismal survival rates. While the genetic features of AML are well understood, the transcriptional programs associated with TP53 dysfunction are less clear, especially across different clinical groups. To fill this gap, we used deep learning on four bulk RNA-seq datasets grouped by p53 expression and a single-cell TARGET-seq dataset classified by TP53 genotype. The bulk data revealed disruptions in chromatin regulation, DNA repair, and immune pathways, with the low-p53 state showing activation of chemokine and interleukin signaling. In the single-cell data, deep learning classifiers successfully distinguished WT from multi-hit TP53-mutant cells, while heterozygous TP53 cells remained transcriptionally similar to WT. Comparing bulk and single-cell results, we found a strong positive correlation between p53-low expression states and TP53-inactive genotypes, linking the expression-based and mutation-based axes of p53 activity. Our study identifies a consistent TP53-related transcriptional signature that connects p53-low expression patterns with underlying TP53 AML.

## Introduction

Acute myeloid leukemia (AML) is a diverse malignancy originating in hematopoietic stem or progenitor cells [1]. Despite risk-based chemotherapy and allogeneic transplantation, the five-year overall survival rate for adults remains at 30% [2]. Genome-wide studies have enhanced classification, yet predicting relapse continues to be difficult.

AML is characterized as a genetically diverse disease that contains multiple rare driver mutations, with no single gene dominating the landscape [3]. Among these genes, alterations in p53 occur in about 7-8% of de novo AML cases. The tumor suppressor gene p53 encodes the protein p53, a transcription factor involved in DNA damage responses, apoptosis, and senescence [4]. p53 consists of four functional domains: (i) an N-terminal transactivation domain rich in phosphorylation sites, (ii) a proline-rich segment, (iii) a sequence-specific DNA-binding domain that can harbor oncogenic mutations, and (iv) a C-terminal oligomerization domain. Recurrent hotspot substitutions such as R175H and R248Q alter the DNA-binding interface, block downstream signaling, and support tumor development. Although less frequent than FLT3-ITD (30%), NPM1 (25-30%), or DNMT3A (20-25%), p53 mutations are associated with the worst overall and event-free survival rates, highlighting the importance of understanding the transcriptional programs that occur with p53 dysfunction.

Consortia such as TCGA, Beat-AML, and the Microarray Innovations in Leukemia (MILE) initiative have documented frequent genetic alterations, but the transcriptional consequences of p53 dysfunction across cohorts remain unclear [5,6]. Earlier studies typically examined one cohort at a time, used straightforward linear tests, and rarely validated their results in an external dataset. New single-cell techniques, including CITE-seq, SHARE-seq, and spatial transcriptomics, can now reveal gene expression in individual cells; however, to our knowledge, no study has combined these data with bulk profiles to develop a single, reliable p53 signature.

In this study, we combine four bulk expression cohorts, including three microarray studies and one RNA-seq dataset [11], and use a deep-learning pipeline to identify transcripts and pathways associated with low and high p53 levels. The resulting p53 signature, which is platform-independent, was then assessed using a single-cell dataset from p53-mutated leukemia patients [12]. Each hematopoietic cell in this dataset is genotyped at allelic resolution and labeled as p53-wild-type, monoallelic (heterozygous) mutant, multihit mutant (M2), or multihit copy-neutral loss-of-heterozygosity (HOM), enabling us to validate and trace the bulk-derived signature back to the specific subpopulations that generated it. This cross-modal approach offers a concise, reproducible marker panel that can support prognosis by serving as a diagnostic tool for early detection of high-risk cases.

## Material and methods

### Single-cell preprocessing

The scRNA-seq data from Rodriguez-Meira et al [12](GSE226340) is represented as a Seurat object containing raw counts for 19,221 cells with pre-applied quality control filters (minimum RNA counts = 2,000 total counts, minimum detected genes = 500, maximum mitochondrial gene expression ratio = 20%). Each sample was normalized separately using SCTransform, with mitochondrial expression ratio regressed out, to correct for differences in sequencing depth. Integration anchors were then identified across all samples, and the data were integrated into a single assay. Principal component analysis was performed on the integrated data, and the first 25 principal components were used to compute a UMAP embedding. Unsupervised clustering with a resolution of 0.5 resulted in 5 distinct clusters.

### Deep learning model for scRNAseq

The single-cell deep learning method has been described in our previous publication [13].

### Gene set construction and overlap analysis

To compare predictors between bulk and single-cell datasets, we first extracted cohort-specific gene sets. For each bulk transcriptome dataset (Metzeler, Verhaak, Haferlach, Vizome), samples were divided into p53-low and p53-high groups using our deep learning framework. Genes were ranked based on their correlation coefficients from the classification models, and the top 2.5% predictors were selected separately for positively and negatively associated genes. This process was repeated for the single-cell models comparing M2 versus preleukemic and HOM versus preleukemic populations. In total, twelve independent gene sets were created: six representing upregulated predictors (top 2.5% positively correlated) and six representing downregulated predictors (top 2.5% negatively correlated).

### GSEA of single-cell models with bulk-derived gene sets

Each of the twelve gene sets was tested using Gene Set Enrichment Analysis (GSEA) against genes preranked by correlation coefficient from the deep learning classifiers of M2 and HOM. Separate analyses were conducted for M2 and HOM, resulting in four enrichment comparisons in total: (i) M2 preranked with up gene sets, (ii) M2 preranked with down gene sets, (iii) HOM preranked with up gene sets, and (iv) HOM preranked with down gene sets.

### Visualization of gene set intersections

For each comparison, overlapping genes between bulk- and single-cell-derived gene sets were visualized using ComplexUpset in R. This method quantifies and displays the overlap among multiple sets, enabling the identification of intersected genes across cohorts and single-cell genotypes.

## Results

### Deep learning-based identification of a pan-leukemic p53-associated gene signature

Mutations in *p53* are linked to one of the poorest survival rates in AML, yet the transcriptional effects of p53 dysfunction are still not well understood. To find consistent gene expression changes related to altered p53 levels, we examined four publicly available transcriptomic datasets with various complex cytogenotypes, including cytogenetically normal AML (CN-AML), chromosomal aberrations-related AML (CA-AML), acute lymphoblastic leukemia (ALL), myelodysplastic syndrome (MDS), myeloproliferative neoplasm (MPN), chronic myeloid leukemia (CML), and chronic lymphocytic leukemia (CLL).

In each cohort independently, Metzeler et al. (CN-AML) [7], Tyner et al. (CN-AML, CA-AML, MDS, MPN) [11], Verhaak et al. (de novo AML) [8] and the MILE study (AML, ALL, CML, CLL, and MDS) [9,10] samples were stratified into p53-high and p53-low groups using unsupervised k-means clustering based on p53 expression (Figure S1A). Deep learning classifiers trained on these stratifications consistently achieved high accuracy on unseen data, ranging from 83.3% to 85.8%, and principal component analysis of the top 5% predictive features revealed robust separation between the two expression states in all datasets (Figure 1A-1D).

**Figure 1.**
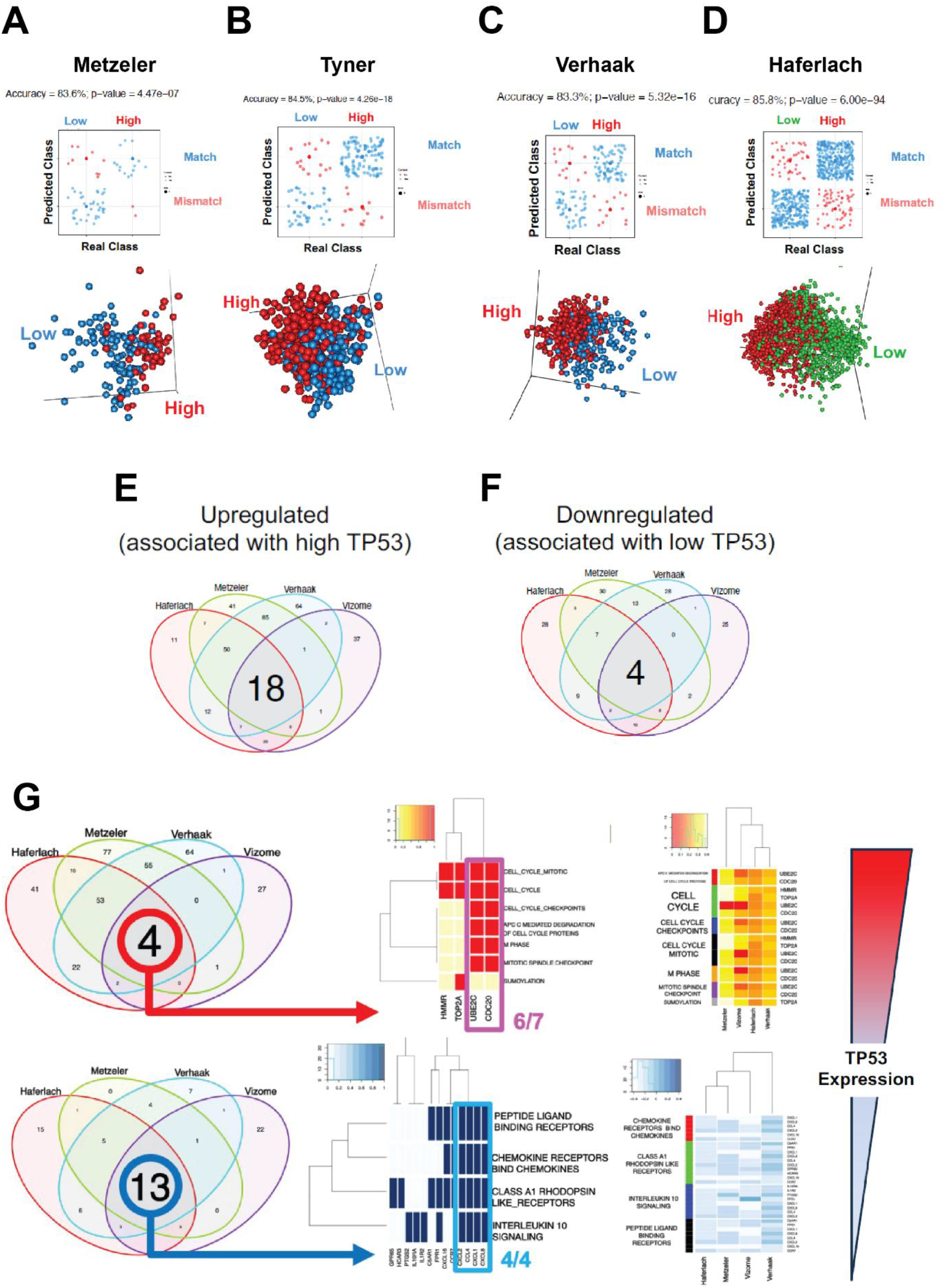
Deep learning-driven identification of a pan-leukemic p53-associated gene signature. **(A-D) Top:** Deep-learning models trained on a subset of **(A)** Metzeler et al., **(B)** Tyner et al., **(C)** Verhaak et al. and **(D)** Haferlach et al. datasets and applied on a subset of unseen data of their corresponding datasets.; Statistical test: Fisher’s exact test. **Bottom**: PCA based on the top 5% of predictors of their corresponding deep-learning models for each of the 4 datasets. **(E, F)** Venn diagram illustrating the overlap of GSEA REACTOME gene sets significantly enriched in either low-p53 or high-p53 expression samples across all four datasets. Gene sets were derived by applying GSEA to the full preranked gene lists from each model, based on deep-learning-derived correlation coefficients. **(G) Left**: Venn diagrams indicating the overlap of core-enriched genes shared among the common gene sets of low p53 and high p53 identified in panel E for each bulk dataset. **Middle**: Binary map indicating presence or absence of those common core-enriched genes for each geneset and dataset. **Right**: Heatmap indicating the running enrichment scores for the intersected genes in their associated pathways across the 4 studies.

For each cohort, predictors from the deep learning models were ranked according to their correlation coefficients and compiled into four pre-ranked gene lists. We first analyzed enriched gene sets within each dataset independently (Figure S1B), then used Panther to identify deregulated biological processes across all datasets in relation to varying p53 levels. Common deregulated processes included genes associated with chromatin modification, DNA repair, metabolism, oxidative stress-induced senescence, and the evasion of oncogene-induced senescence (Figure S1C and Table S1).

Next, using REACTOME-based GSEA, we identified intersecting enriched gene sets across all four datasets (Figure 1E and 1F). In the p53-high groups, 18 gene sets were consistently enriched, with four core genes appearing across all four data sets, mainly related to cell cycle control and SUMOylation (Figure 1E and 1G). In contrast, for the p53-low groups, only four gene sets were shared across datasets, yet they included 13 common core genes (Figure 1F and 1G). These genes are involved in peptide signaling, rhodopsin-like receptor pathways, and chemokine and interleukin signaling (Figure 1G).

Taken together, deep-learning analysis of four independent cohorts identified reproducible gene signatures linked to *p53* expression. These signatures remained consistent across different technologies, disease subtypes, and geographic regions. Importantly, this analysis uncovered a new association between low p53 expression and the deregulation of chemokine and interleukin signaling pathways, indicating an immune-related transcriptional state connected to reduced p53 levels.

### Deep learning detects p53-mutant states at single-cell resolution

We next analyzed the TARGET-seq dataset from Rodriguez-Meira et al. [12], which provides per-cell transcriptomes along with matched TP53 genotypes. Cells were assigned to four groups: wild-type (WT), monoallelic mutant (HET), multi-hit biallelic mutant (M2), and multi-hit copy-neutral loss of heterozygosity (HOM). This framework allows us to test whether the bulk p53-associated signature maps to specific genotype-defined subpopulations.

We first calculated the joint UMAP from the single-cell data. Unsupervised clustering grouped the cells into five transcriptionally distinct clusters (Figure S2A, S2B). Importantly, visualizing the cells in the UMAP based on sequencing instrument, patient, tissue of origin, or their TP53 genotype showed widespread mixing without isolated clusters, indicating successful integration and that batch effects were managed (Figure 2A, S2C).

**Figure 2.**
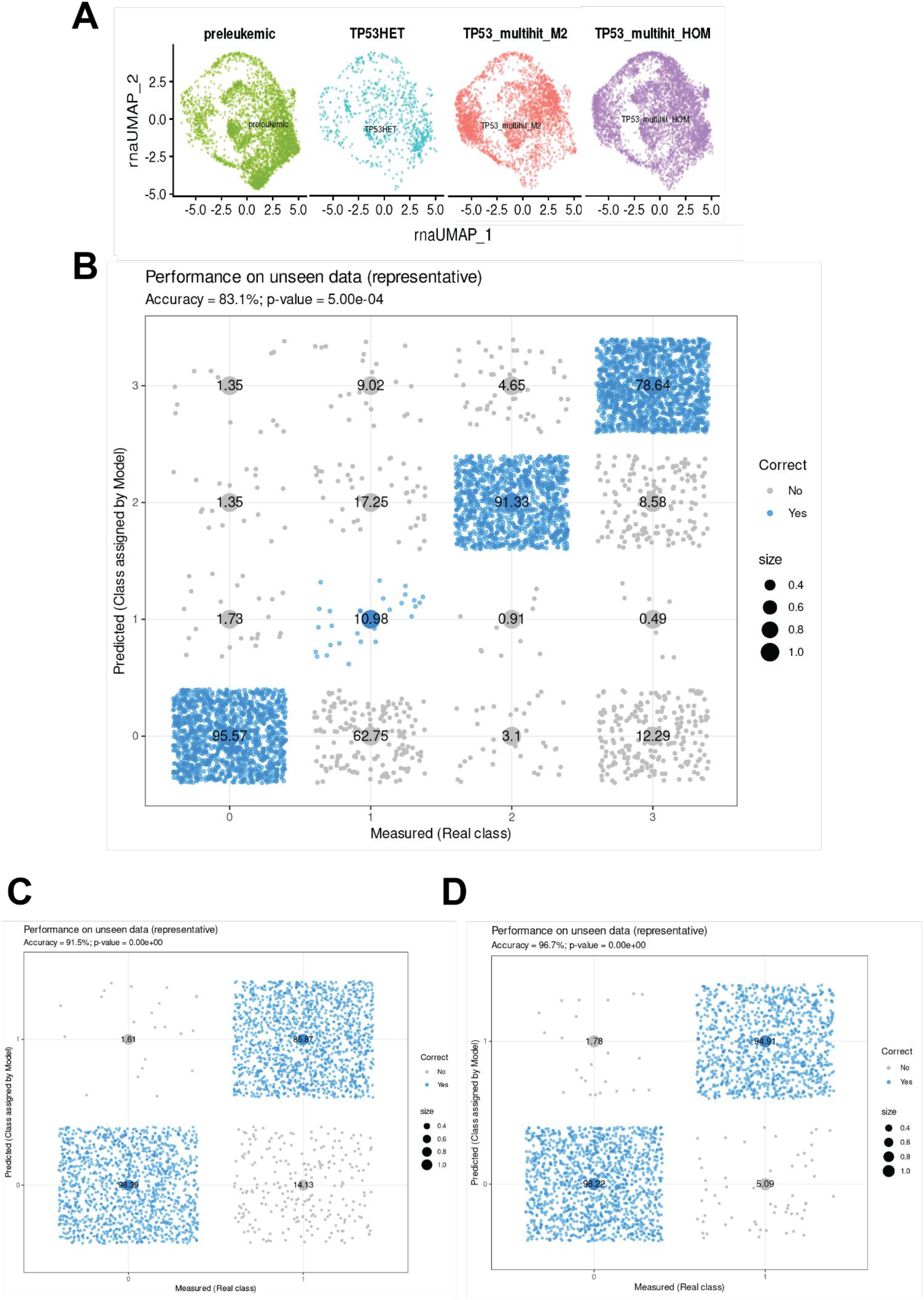
Deep learning identifies p53-mutant states at single-cell resolution. **(A)** UMAP colored by p53 genotype, displayed per genotype. **(B)** Confusion matrix on unseen data of a four-class deep learning model trained to classify WT, HET, M2, and HOM cells.; Statistical test: Fisher’s exact test. **(C, D)** Confusion matrices on unseen data of binary deep learning models: **(C)** WT vs HOM and **(D)** WT vs M2.; Statistical test: Fisher’s exact test.

We then explored whether the full transcriptome could determine TP53 genotype and uncover hidden relationships at the single-cell level. A four-class deep learning classifier trained to predict WT, HET, M2, or HOM achieved 83.1% accuracy across all cells (Figure 2B); however, class-specific errors varied. WT, M2, and HOM were clearly separated, while HET cells were often misclassified as WT. This pattern suggests that heterozygous TP53 mutations induce a weaker transcriptional shift compared to M2 or HOM.

To verify these trends at the sample level and reduce single-cell noise, we pseudo-bulked cells by patient within each genotype and trained pairwise random-forest models (Table S2-S4). Performance evaluation replicated the single-cell results: WT vs HOM yielded an out-of-bag (OOB) misclassification rate of only 10%; WT vs M2 yielded 0 OOB error, while WT vs HET yielded 40%. Finally, given our focus on multi-hit p53-mutant states, we trained two binary deep-learning classifiers on the single-cell level, comparing WT with HOM and WT with M2. The models achieved 91.5% and 96.7% accuracy, respectively (Figure 2C, 2D), demonstrating clear separability of their transcriptomes based on TP53 genotype.

Taken together, the single-cell deep learning classifiers show a clear separation of WT from multi-hit p53-mutant genotypes (M2, HOM) but not from HET. This aligns with the p53 dosage theory, since HET retains one functional allele and remains transcriptionally similar to WT at baseline. As a result, clear separation only occurs when both alleles are altered (M2 or HOM).

### Low P53 expression transcriptome correlates with multi-hit TP53 mutant profiles

After assembling the transcriptome differences associated with TP53 genotype from both bulk and single-cell analyses, we examined the correlation among the four bulk-derived p53 datasets (Metzeler, Vizome, Verhaak, Haferlach) and the single-cell dataset (preleukemic and M2/HOM). As shown in Figure 3A, the bulk datasets were strongly correlated with each other and also positively correlated with the M2 group. In contrast, the preleukemic WT population showed negative correlation with each bulk p53-low dataset and with M2. Repeating the analysis with HOM instead of M2 produced the same pattern (Figure 3B). Therefore, our analysis emphasizes the positive correlation of transcriptomes between low p53 expression, derived from bulk analysis, and multi-hit TP53, derived from scRNAseq analysis.

**Figure 3.**
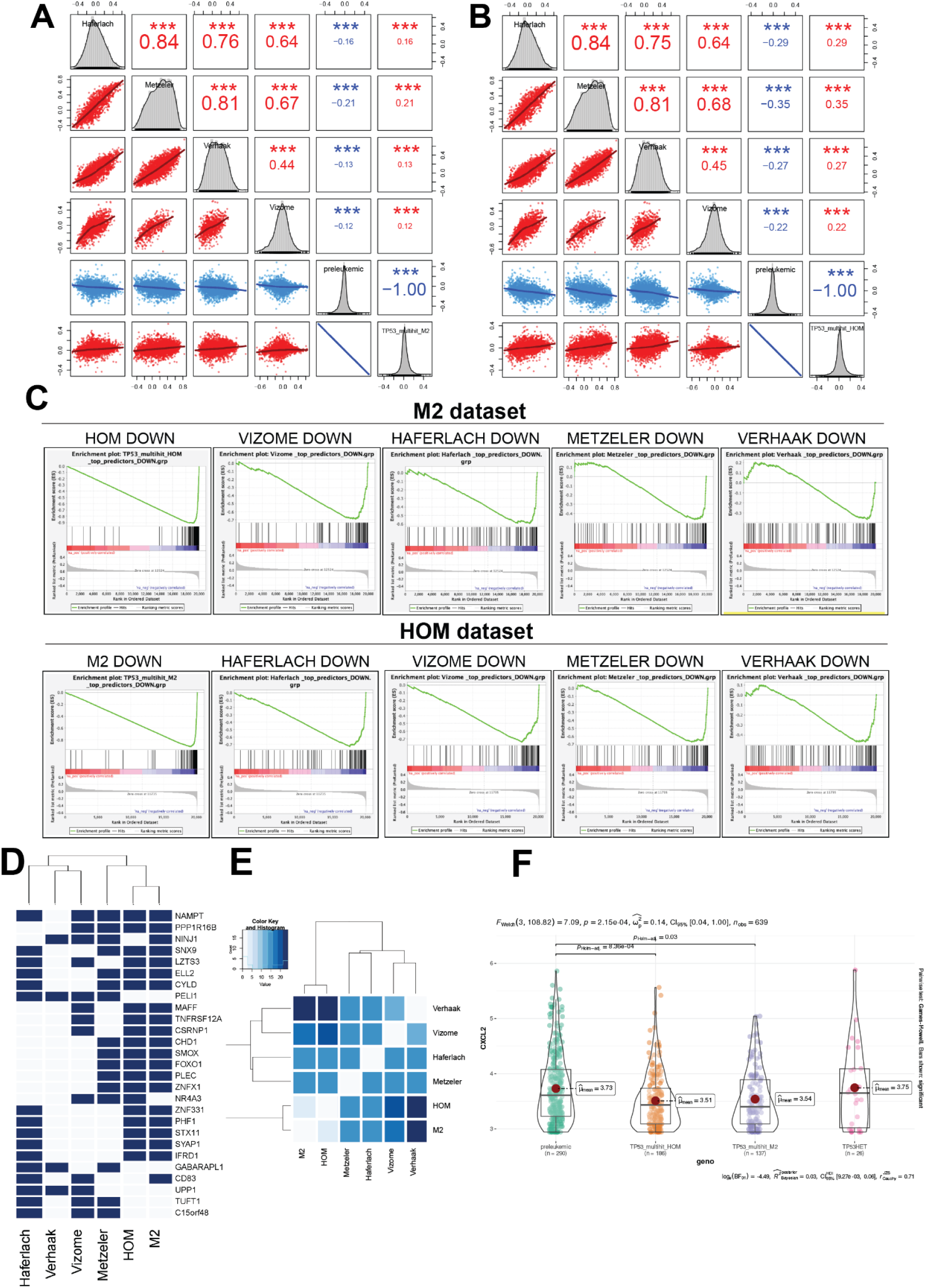
CXCL2 emerges as a shared marker of low-p53 bulk and p53-mutant single cells. **(A, B)** Correlation matrices of bulk-derived p53 signatures (Haferlach, Metzeler, Verhaak, Vizome) with **(A)** M2 and **(B)** HOM single-cell populations. Upper panels display Pearson correlation coefficients with significance levels; lower panels show scatter plots. **(C)** Gene set enrichment analysis (GSEA) plots showing depletion of bulk-derived low-p53 downregulated genes as genesets in M2 and HOM single-cell preranked genes. **(D)** Binary presence-absence map showing overlap of downregulated genes identified across bulk p53-low (Haferlach, Verhaak, Vizome, Metzeler) and multihit single-cell (M2, HOM) datasets. Dark blue bars indicate genes present in each dataset; clustering above reflects similarity in gene occurrence patterns among populations. **(E)** Hamming distance heatmap showing the degree of similarity among core downregulated genes identified across six datasets (Haferlach, Verhaak, Vizome, Metzeler, HOM, and M2). **(F)** Violin plots of CXCL2 expression in single-cell populations (preleukemic, HOM, M2, HET). Box plots indicate interquartile range with medians. Red points mark group means. Pairwise comparisons were performed using Games-Howell tests.

Given the consistent transcriptional correlation between the low-p53 bulk and the M2/HOM populations, we next examined whether they shared genes showing coordinated expression changes across both analyses. Therefore, we compiled gene sets consisting of the top downregulated genes from each of the low-p53 bulk datasets, as well as from the M2 and HOM single-cell datasets, and used these for gene set enrichment analysis (GSEA). We then performed GSEA on the pre-ranked gene lists from H2 and HOM against these compiled gene sets. All gene sets showed significant depletion in both M2 and HOM datasets, indicating that the genes most downregulated in low-p53 bulk samples were also consistently suppressed in these TP53-mutant populations (Figure 3C). These findings reinforce the strong agreement between the transcriptomic profiles of low-p53 bulk samples and multi-hit TP53-inactive single-cell populations.

To identify the core genes consistently depleted across datasets, we intersected the six gene sets derived from the M2 and HOM GSEA results (Figure S3A–D). Focusing on genes present in at least four of these sets, we identified a recurring group of downregulated genes, including NAMPT, PPP1R16B, NINJ1, DNX9, LZTS2, ELL2, and CYLD, many of which have well-established roles in cancer biology (Figure 3D). A complementary Hamming distance analysis further confirmed the extent of overlap among these gene sets, emphasizing their shared depletion patterns (Figure 3E).

Finally, based on the low-p53-associated REACTOME signatures identified in the bulk data, we examined whether specific genes from these pathways also appeared as key features in the single-cell models. The bulk analysis revealed 13 core genes that were consistently enriched across cohorts. To check for overlap, we intersected these with the top 2.5% predictors from our deep learning models comparing M2 and HOM to preleukemic cells. Notably, CXCL2 was also identified in the single-cell analysis, ranking among the top predictors for both M2 and HOM and also present in the low-p53 bulk core set (Figure 3F).

## Discussion

Acute myeloid leukemia (AML) is a genetically diverse cancer, with TP53 changes linked to dismal outcomes. Although several studies have looked at TP53-mutated AML individually, most have focused on one or two datasets at a time, which limits broader applicability[14–19]. In this study, we combined four independent bulk transcriptome datasets (Metzeler, Vizome, Verhaak, and MILE) with a single-cell TARGET-seq dataset that included matched TP53 genotypes. This method enabled us to identify transcriptional signatures related to TP53 status across various platforms.

To analyze TP53-associated transcriptional changes without bias across bulk transcriptomic cohorts, we used unsupervised k-means clustering to group samples based on TP53 expression levels. Despite variations in cohort size, patient populations, sequencing platforms, and disease composition, our deep learning models maintained consistent accuracies (83–86%), demonstrating strong performance across datasets. To our knowledge, no previous study has used deep learning to identify predictive signatures for p53 stratification across multiple independent AML cohorts. This method ensured reliable classification and allowed us to extract gene-level predictors linked to p53 levels. Importantly, no prior research has integrated this many independent bulk transcriptomic datasets to develop a unified TP53-associated signature, underscoring the novelty of our multi-cohort approach.

Panther analysis identified key biological processes, including chromatin modification, DNA repair, and cell cycle regulation, as among the most deregulated between the p53-low and p53-high expression groups. These findings reinforce the well-established role of p53 in maintaining genomic integrity through DNA repair and cell cycle regulation. Similar pathway enrichment, as observed in previous transcriptomic studies of TP53-mutated AML and MDS, suggests that p53 alterations disrupt cell cycle and chromatin organization [20,21].

Importantly, p53-low samples showed enrichment in chemokine signaling pathways, including genes such as CXCL1, CXCL2, CXCL8, and CCL4. This immune-related state of p53-low groups was found in all four datasets. Previous studies have reported similar trends, although usually within single cohorts. For example, TP53-mutated AML and MDS were reported to display inflammatory and interferon-related gene signatures, including CXCL1, CXCL2, and CXCL8 [16]. Similarly, it has been shown that TP53-mutant cases are linked to immune infiltration and IL2–STAT5 pathway activation using the TCGA and BeatAML cohorts [17]. Conversely, a recent study identified immune suppression and decreased innate and adaptive immune responses in TP53-dysfunctional MDS [19]. These differences likely result from variations in immune composition, while our analysis focuses on transcriptional pathways rather than specific immune cell populations. Overall, our multi-cohort analysis indicates that p53 dysfunction is associated with deregulation of immune signaling and cell cycle pathways, pointing to a broader disruption of inflammatory transcriptional control.

Analysis of the single-cell TARGET-seq dataset provided further insight into how TP53 genotype affects transcriptional states in AML. Using matched TP53 genotypes, our deep learning models accurately distinguished TP53-inactive cells (M2 and HOM) from WT cells with 91–97% accuracy. In contrast, heterozygous mutants were often classified as WT, indicating that monoallelic TP53 loss alone does not cause significant transcriptomic changes. These findings emphasize a dosage-dependent effect of p53, where transcriptional reprogramming only occurs after complete loss of p53 function. This study is the first to show that heterozygous TP53 inactivation does not produce a distinct transcriptomic profile in AML. These results align with clinical data indicating limited prognostic impact of monoallelic TP53 mutations compared with wild-type, highlighting the importance of allelic dosage in myeloid neoplasias. [22,23].

Correlation analysis between bulk and single-cell datasets revealed that TP53-mutant single-cell profiles were linked to p53-low expression groups from bulk data, while WT cells showed an opposite pattern. This suggests that the transcriptional programs observed in bulk analysis mirror the TP53 genotype at the cellular level, connecting expression-based and mutation-based axes of p53 activity. Gene set enrichment analysis further confirmed this relationship by identifying common predictors between datasets, including immune-related and metabolic factors such as CXCL2 and NAMPT. The correspondence between bulk p53-low signatures and biallelic TP53-mutant single cells emphasizes that loss of p53 function drives a consistent transcriptional program detectable in both bulk and single-cell analyses. Overall, our study goes beyond previous transcriptomic or mutational research by linking TP53 dosage to transcriptional states across bulk and single-cell data.

In conclusion, our study combines bulk transcriptomic and single-cell data to define transcriptional states linked with TP53 activity in AML. Using deep learning across multiple independent cohorts, we identified consistent TP53-related signatures that are reproducible across disease subtypes and various experimental platforms. This is the first study to merge deep learning with cross-resolution transcriptomic integration, directly connecting the expression-based and mutation-based axes of p53 activity. We offer a comprehensive view of how TP53 inactivation influences downstream transcriptional programs in AML. Future research incorporating proteomic and spatial analyses will be essential to understand how these transcriptional programs affect the leukemic microenvironment and responses to therapy.

## Data availability statement

This study reanalyzes previously published datasets that are publicly available through the studies cited in the manuscript. No new data were generated or deposited.

## Acknowledgments

We acknowledge the funding sources: European Research Council (ERC) grant 101002453 (MCF) and LaCaixa Health Research grant HR20-00800 (MCF & MAM).

## Authorship contributions

Conceptualization: MAM, MCF

Methodology: MAM, MT, RA, SM-V, AF-P

Investigation: MT, RA, SM-V, AF-P

Visualization: MAM, MT

Funding acquisition: MAM, MCF

Project administration: MAM, MCF

Supervision: MAM, MCF

Writing – original draft: MT, RA, MAM, MCF

Writing – review & editing: MAM, MCF

## References

1. Wachter F, Pikman Y. Pathophysiology of Acute Myeloid Leukemia. Acta Haematol. 2024;147:229– 46. 10.1159/000536152

2. Shimony S, Stahl M, Stone RM. Acute Myeloid Leukemia: 2025 Update on Diagnosis, Risk-Stratification, and Management. Am J Hematol. 2025;100:860–91. 10.1002/ajh.27625

3. Molica M, Mazzone C, Niscola P, de Fabritiis P. TP53 Mutations in Acute Myeloid Leukemia: Still a Daunting Challenge? Front Oncol. 2020;10:610820. 10.3389/fonc.2020.610820

4. Vousden KH, Lane DP. p53 in health and disease. Nat Rev Mol Cell Biol. Nature Publishing Group; 2007;8:275–83. 10.1038/nrm2147

5. Haferlach T, Kohlmann A, Wieczorek L, Basso G, Kronnie GT, Béné M-C, et al. Clinical utility of microarray-based gene expression profiling in the diagnosis and subclassification of leukemia: report from the International Microarray Innovations in Leukemia Study Group. J Clin Oncol Off J Am Soc Clin Oncol. 2010;28:2529–37. 10.1200/JCO.2009.23.4732

6. Tyner JW, Tognon CE, Bottomly D, Wilmot B, Kurtz SE, Savage SL, et al. Functional Genomic Landscape of Acute Myeloid Leukemia. Nature. 2018;562:526–31. 10.1038/s41586-018-0623-z

7. Metzeler KH, Hummel M, Bloomfield CD, Spiekermann K, Braess J, Sauerland M-C, et al. An 86-probe-set gene-expression signature predicts survival in cytogenetically normal acute myeloid leukemia. Blood. 2008;112:4193–201. 10.1182/blood-2008-02-134411

8. Verhaak RGW, Wouters BJ, Erpelinck CAJ, Abbas S, Beverloo HB, Lugthart S, et al. Prediction of molecular subtypes in acute myeloid leukemia based on gene expression profiling. Haematologica. 2009;94:131–4. 10.3324/haematol.13299

9. Kohlmann A, Kipps TJ, Rassenti LZ, Downing JR, Shurtleff SA, Mills KI, et al. An international standardization programme towards the application of gene expression profiling in routine leukaemia diagnostics: the Microarray Innovations in LEukemia study prephase. Br J Haematol. Wiley; 2008;142:802–7. 10.1111/j.1365-2141.2008.07261.x

10. Haferlach T, Kohlmann A, Wieczorek L, Basso G, Kronnie GT, Béné M-C, et al. Clinical utility of microarray-based gene expression profiling in the diagnosis and subclassification of leukemia: report from the International Microarray Innovations in Leukemia Study Group. J Clin Oncol Off J Am Soc Clin Oncol. 2010;28:2529–37. 10.1200/JCO.2009.23.4732

11. Tyner JW, Tognon CE, Bottomly D, Wilmot B, Kurtz SE, Savage SL, et al. Functional genomic landscape of acute myeloid leukaemia. Nature. Nature Publishing Group; 2018;562:526–31. 10.1038/s41586-018-0623-z

12. Rodriguez-Meira A, Norfo R, Wen S, Chédeville AL, Rahman H, O’Sullivan J, et al. Single-cell multiomics identifies chronic inflammation as a driver of TP53-mutant leukemic evolution. Nat Genet. 2023;55:1531–41. 10.1038/s41588-023-01480-1

13. Hageb A, Thalheim T, Nattamai KJ, Möhrle B, Saçma M, Sakk V, et al. Reduced adhesion of aged intestinal stem cells contributes to an accelerated clonal drift. Life Sci Alliance. 2022;5:e202201408. 10.26508/lsa.202201408

14. Tashakori M, Kadia T, Loghavi S, Daver N, Kanagal-Shamanna R, Pierce S, et al. TP53 copy number and protein expression inform mutation status across risk categories in acute myeloid leukemia. Blood. 2022;140:58–72. 10.1182/blood.2021013983

15. Xie J, Chen K, Han H, Dong Q, Wang W. Establishment of tumor protein p53 mutation-based prognostic signatures for acute myeloid leukemia. Curr Res Transl Med. 2022;70:103347. 10.1016/j.retram.2022.103347

16. Vadakekolathu J, Lai C, Reeder S, Church SE, Hood T, Lourdusamy A, et al. TP53 abnormalities correlate with immune infiltration and associate with response to flotetuzumab immunotherapy in AML. Blood Adv. 2020;4:5011–24. 10.1182/bloodadvances.2020002512

17. Wen X, Xu Z, Jin Y, Xia P, Ma J, Qian W, et al. Association Analyses of TP53 Mutation With Prognosis, Tumor Mutational Burden, and Immunological Features in Acute Myeloid Leukemia. Front Immunol. 2021;12:717527. 10.3389/fimmu.2021.717527

18. Cucchi DGJ, Bachas C, Klein K, Huttenhuis S, Zwaan CM, Ossenkoppele GJ, et al. TP53 mutations and relevance of expression of TP53 pathway genes in paediatric acute myeloid leukaemia. Br J Haematol. 2020;188:736–9. 10.1111/bjh.16229

19. Zampini M, Riva E, Lanino L, Sauta E, Antunes Dos Reis R, Ejarque RMA, et al. Characterization and Clinical Implications of p53 Dysfunction in Patients With Myelodysplastic Syndromes. J Clin Oncol. Wolters Kluwer; 2025;43:2069–83. 10.1200/JCO-24-02394

20. Shahzad M, Amin MK, Daver NG, Shah MV, Hiwase D, Arber DA, et al. What have we learned about TP53-mutated acute myeloid leukemia? Blood Cancer J. Nature Publishing Group; 2024;14:202. 10.1038/s41408-024-01186-5

21. Cumbo C, Tota G, Anelli L, Zagaria A, Specchia G, Albano F. TP53 in Myelodysplastic Syndromes: Recent Biological and Clinical Findings. Int J Mol Sci. Multidisciplinary Digital Publishing Institute; 2020;21:3432. 10.3390/ijms21103432

22. Bahaj W, Kewan T, Gurnari C, Durmaz A, Ponvilawan B, Pandit I, et al. Novel scheme for defining the clinical implications of TP53 mutations in myeloid neoplasia. J Hematol OncolJ Hematol Oncol. 2023;16:91. 10.1186/s13045-023-01480-y

23. Bernard E, Nannya Y, Hasserjian RP, Devlin SM, Tuechler H, Medina-Martinez JS, et al. Implications of TP53 allelic state for genome stability, clinical presentation and outcomes in myelodysplastic syndromes. Nat Med. Nature Publishing Group; 2020;26:1549–56. 10.1038/s41591-020-1008-z

