## Supplementary Files (PDF) for "Deep Learning links TP53 genotype to expression-defined transcriptional program in Acute Myeloid Leukemia"

### Figure S1

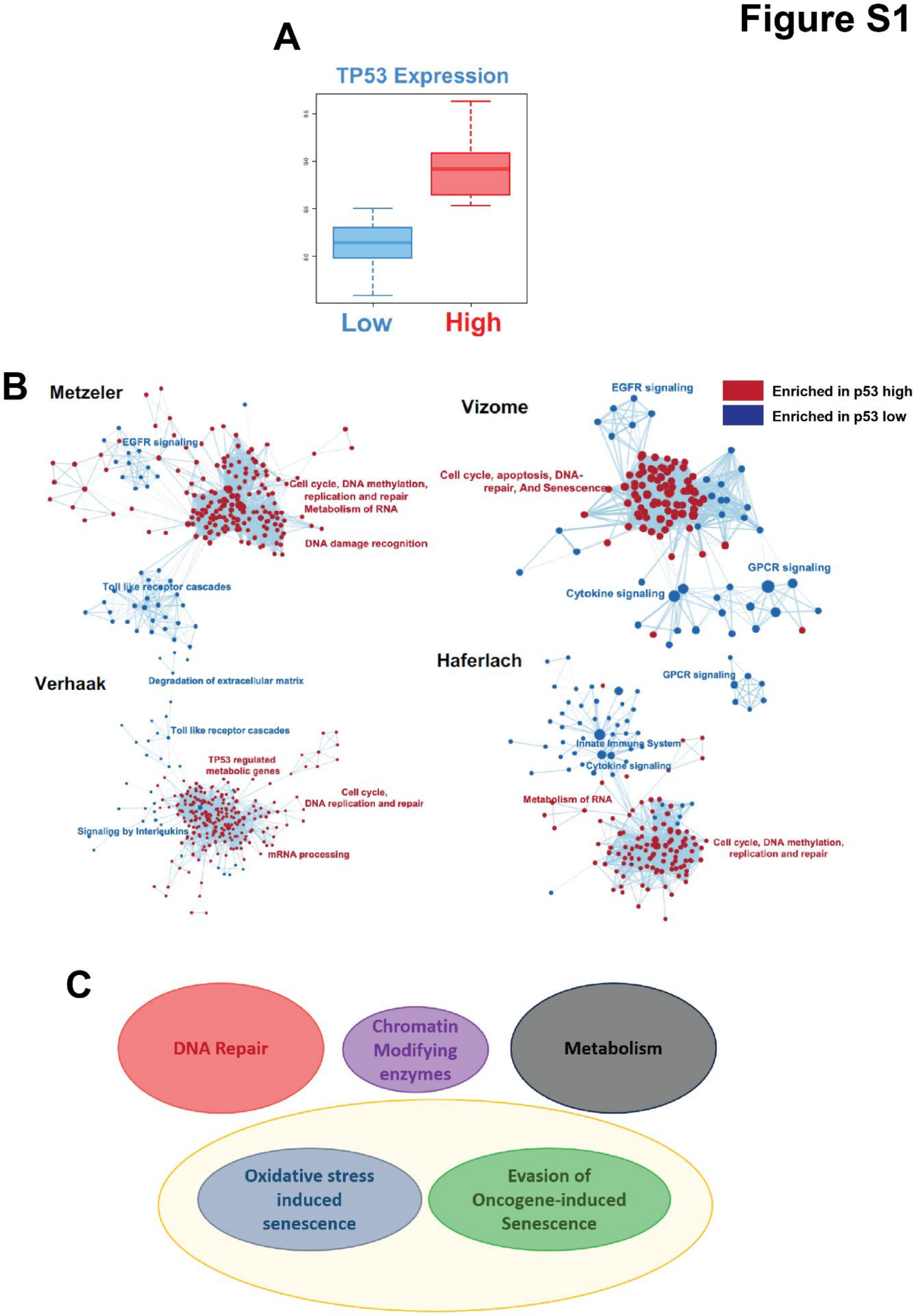
A
TP53 Expression
Low
High
B
Metzeler
EGFR signaling
Cell cycle, DNA methylation, replication and repair
Metabolism of RNA
DNA damage recognition
Toll like receptor cascades
Degradation of extracellular matrix
Vizome
EGFR signaling
Cell cycle, apoptosis, DNA-repair, And Senescence
GPCR signaling
Cytokine signaling
Enriched in p53 high
Enriched in p53 low
Verhaak
Toll like receptor cascades
TP53 regulated metabolic genes
Cell cycle, DNA replication and repair
Signaligg by Interleukins
mRNA processing
Haferlach
GPCR signaling
Innate Immune System
Cytokine signaling
Metabolism of RNA
Cell cycle, DNA methylation, replication and repair
C
DNA Repair
Chromatin Modifying enzymes
Metabolism
Oxidative stress induced senescence
Evasion of Oncogene-induced Senescence

**Figure S1: (A)** k-means unsupervised clustering of Metzeler et al dataset visualized as an example of how p53 low and p53 high populations were classified according to the p53 level of each sample. **(B)** Enrichment map of significantly enriched GSEA REACTOME genesets of p53 low or p53 high for each of the 4 datasets independently. Red nodes signify enrichment in the p53-high group and blue nodes signify enrichment in the p53-low group. **(C)** Visualization of Panther database results indicating deregulated processes concerning different p53 levels across all 4 datasets.

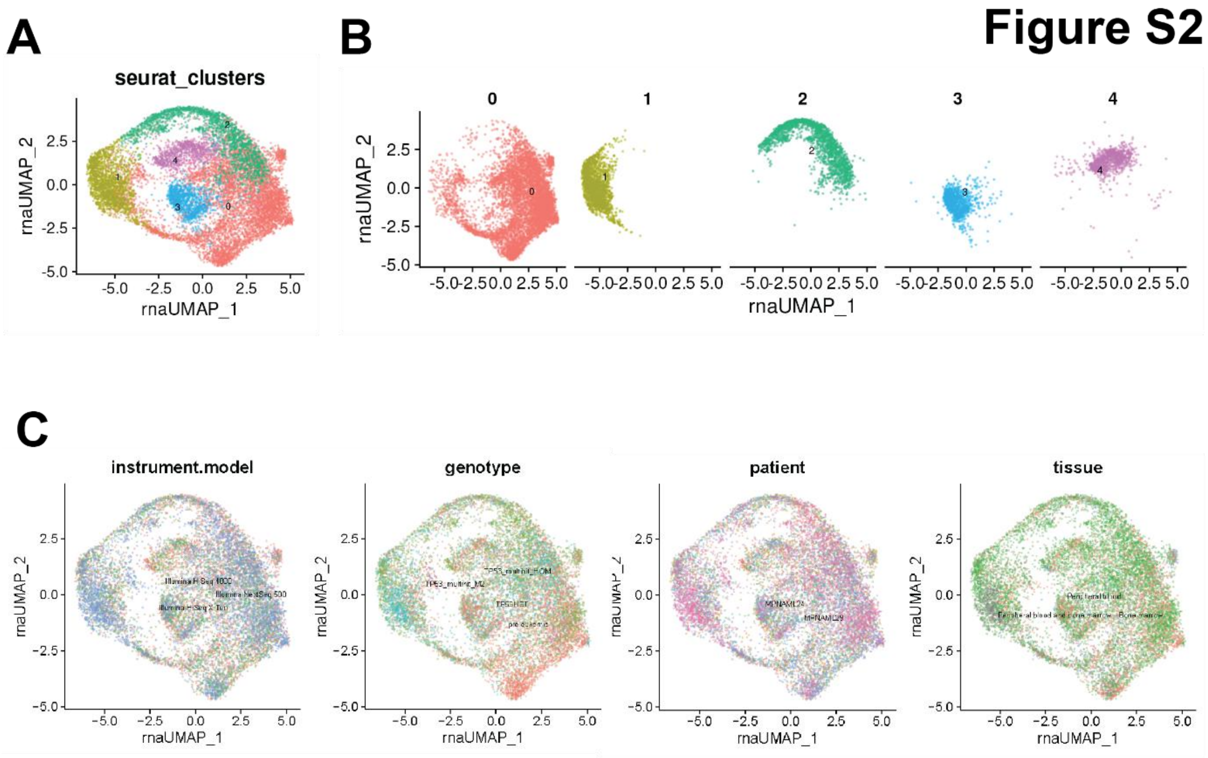


**Figure S2**: **(A, B)** Unsupervised clustering at resolution 0.5 identifying five populations with **(A)** all clusters combined and **(B)** each cluster shown separately. **(C)** UMAP colored by sequencing platform, patient, tissue, and genotype.

**Figure S3**

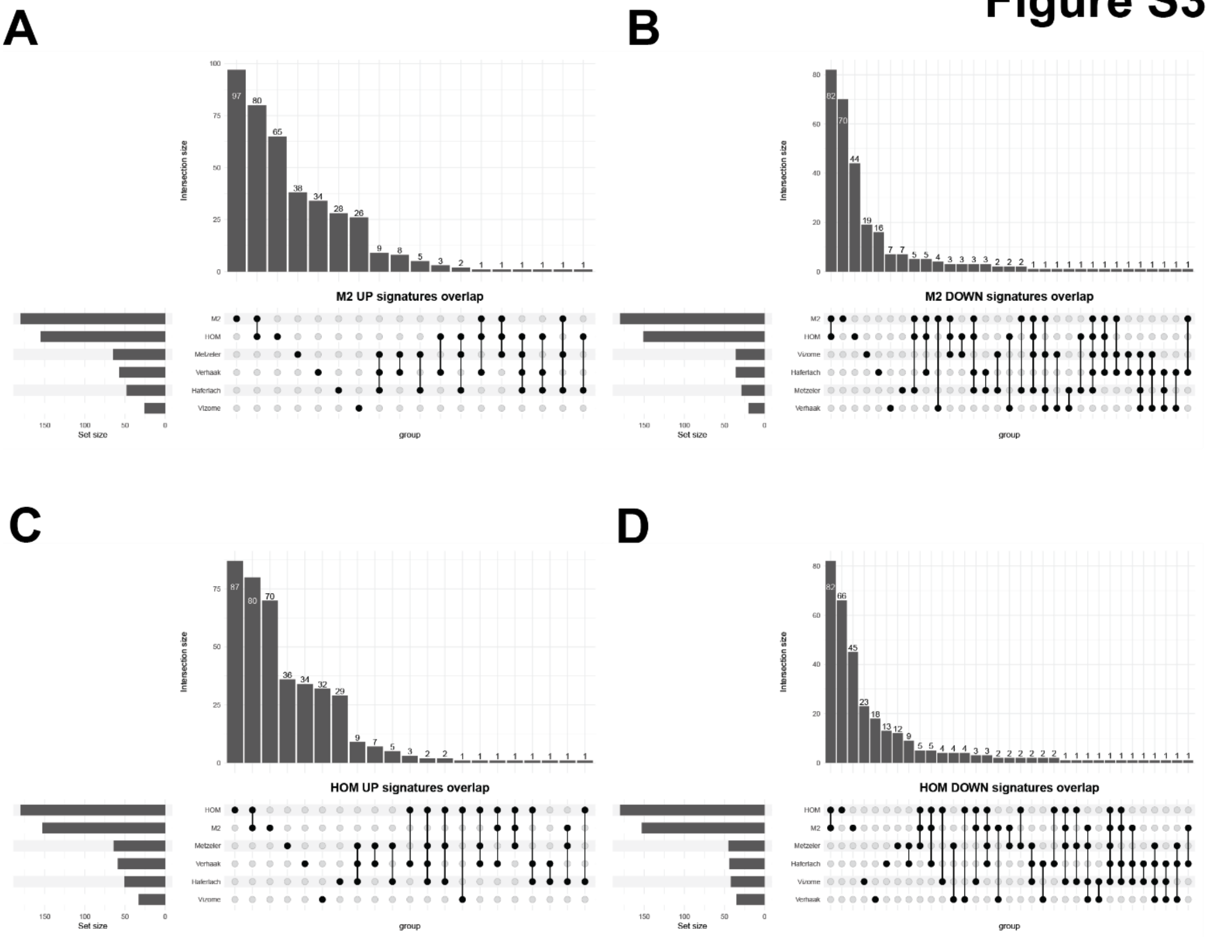


**Figure S3**: **(A-D)** Intersections of gene sets derived from the top 2.5% predictors of four bulk cohorts and two single-cell comparisons (M2 vs preleukemic, HOM vs preleukemic). Up- and downregulated predictors were separated, yielding six up and six down gene sets. These sets were tested by GSEA against preranked lists from the deep learning models of M2 and HOM. **(A)** M2 preranked with up sets. **(B)** M2 preranked with down sets. **(C)** HOM preranked with up sets. **(D)** HOM preranked with down sets.

| Pathway | Reactome ID |
|---|---|
| Purine ribonucleoside monophosphate biosynthesis | R-HSA-73817 |
| Nucleobase biosynthesis | R-HSA-8956320 |
| Metabolism of nucleotides | R-HSA-15869 |
| DNA Damage Recognition in GG-NER | R-HSA-5696394 |
| Evasion of Oxidative Stress Induced Senescence Due to Defective p16INK4A binding to CDK4 | R-HSA-9632697 |
| Evasion of Oncogene Induced Senescence Due to Defective p16INK4A binding to CDK4 | R-HSA-9630791 |
| Defective Mismatch Repair Associated With MSH6 | R-HSA-5632968 |
| Defective Mismatch Repair Associated With MSH3 | R-HSA-5632927 |
| Evasion of Oxidative Stress Induced Senescence Due to p16INK4A Defects | R-HSA-9632693 |
| Evasion of Oncogene Induced Senescence Due to Defective p16INK4A binding to CDK4 and CDK6 | R-HSA-9630794 |
| Evasion of Oncogene Induced Senescence Due to p16INK4A Defects | R-HSA-9630750 |
| Diseases of Cellular Senescence | R-HSA-9630747 |
| Evasion of Oxidative Stress Induced Senescence Due to Defective p16INK4A binding to CDK4 and CDK6 | R-HSA-9632700 |
| Defective Mismatch Repair Associated With MSH2 | R-HSA-5632928 |
| Chromatin modifying enzymes | R-HSA-3247509 |
| Chromatin organization | R-HSA-4839726 |
| Global Genome Nucleotide Excision Repair (GG-NER) | R-HSA-5696399 |
| Diseases of Mismatch Repair (MMR) | R-HSA-5423599 |
| Beta oxidation of octanoyl-CoA to hexanoyl-CoA | R-HSA-77348 |

**Table S1**: Reactome sets that were examined across all 4 bulk transcriptome cohorts, including their Reactome IDs**.**

| | | | |
|---|---|---|---|
| Sample size | 10 | | |
| Frequency of class labels | 5 | 5 | |
| Number of trees | 500 | | |
| Forest terminal node size | 1 | | |
| Average no. of terminal nodes | 2 | | |
| No. of variables tried at each split | 148 | | |
| Total no. of variables | 21648 | | |
| Resampling used to grow trees | swor | | |
| Resample size used to grow trees | 6 | | |
| Analysis | RF-C | | |
| Family | Class | | |
| Splitting rule | gini *random* | | |
| Number of random split points | 10 | | |
| Imbalanced ratio | 1 | | |
| (OOB) Brier score | 0.18192487 | | |
| (OOB) Normalized Brier score | 0.72769949 | | |
| (OOB) AUC | 1 | | |
| (OOB) Log-loss | 0.55303108 | | |
| (OOB) PR-AUC | 1 | | |
| (OOB) G-mean | 0.89442719 | | |
| (OOB) Requested performance error | 0.1 | 0.2 | 0 |
| **Confusion matrix:** | | | |
| | predicted preleukemic | predicted TP53-multihit-M2 | class.error |
| observed preleukemic | 4 | 1 | 0.2 |
| observed TP53-multihit-M2 | 0 | 5 | 0 |
| **(OOB) Misclassification rate** | 0.1 | | |

**Table S2**: Random forest performance distinguishing preleukemic vs TP53-multihit M2 (OOB metrics and confusion matrix).

| | | | |
|---|---|---|---|
| Sample size | 10 | | |
| Frequency of class labels | 5 | 5 | |
| Number of trees | 500 | | |
| Forest terminal node size | 1 | | |
| Average no. of terminal nodes | 2 | | |
| No. of variables tried at each split | 148 | | |
| Total no. of variables | 21648 | | |
| Resampling used to grow trees | swor | | |
| Resample size used to grow trees | 6 | | |
| Analysis | RF-C | | |
| Family | Class | | |
| Splitting rule | gini *random* | | |
| Number of random split points | 10 | | |
| Imbalanced ratio | 1 | | |
| (OOB) Brier score | 0.18168593 | | |
| (OOB) Normalized Brier score | 0.72674373 | | |
| (OOB) AUC | 1 | | |
| (OOB) Log-loss | 0.55370164 | | |
| (OOB) PR-AUC | 1 | | |
| (OOB) G-mean | 1 | | |
| (OOB) Requested performance error | 0 | 0 | 0 |
| **Confusion matrix:** | | | |
| | predicted preleukemic | predicted TP53-multihit-HOM | class.error |
| observed preleukemic | 5 | 0 | 0 |
| observed TP53-multihit-HOM | 0 | 5 | 0 |
| **(OOB) Misclassification rate** | 0 | | |

**Table S3**: Random forest performance distinguishing preleukemic vs TP53-multihit HOM (OOB metrics and confusion matrix).

| | | | |
|---|---|---|---|
| Sample size | 10 | | |
| Frequency of class labels | 5 | 5 | |
| Number of trees | 500 | | |
| Forest terminal node size | 1 | | |
| Average no. of terminal nodes | 2 | | |
| No. of variables tried at each split | 148 | | |
| Total no. of variables | 21648 | | |
| Resampling used to grow trees | swor | | |
| Resample size used to grow trees | 6 | | |
| Analysis | RF-C | | |
| Family | Class | | |
| Splitting rule | gini *random* | | |
| Number of random split points | 10 | | |
| Imbalanced ratio | 1 | | |
| (OOB) Brier score | 0.228732 | | |
| (OOB) Normalized Brier score | 0.914928 | | |
| (OOB) AUC | 0.76 | | |
| (OOB) Log-loss | 0.65010808 | | |
| (OOB) PR-AUC | 0.78941613 | | |
| (OOB) G-mean | 0.6 | | |
| (OOB) Requested performance error | 0.4 | 0.4 | 0.4 |
| **Confusion matrix:** | | | |
| | predicted preleukemic | predicted HET | class.error |
| observed preleukemic | 3 | 2 | 0.4 |
| observed TP53-multihit-HOM | 2 | 3 | 0.4 |
| **(OOB) Misclassification rate** | 0.4 | | |

**Table S4**: Random forest performance distinguishing preleukemic vs TP53-HET (OOB metrics and confusion matrix).
